# FRESH™ 3D Bioprinted Cardiac Tissue, a Bioengineered Platform for in vitro Toxicology and Pharmacology

**DOI:** 10.1101/2023.03.13.528447

**Authors:** Samuel Finkel, Shannon Sweet, Tyler Locke, Sydney Smith, Zhefan Wang, Christopher Sandini, John P. Imredy, Yufang He, Marc Durante, Armando Lagrutta, Adam Feinberg, Andrew Lee

**Author notes:** **Correspondence:** Andrew Lee.

## Abstract

There is critical need for a predictive model of human cardiac physiology in the drug development process for assessment of compound toxicology and pharmacology. In vitro two-dimensional monolayer culture of cardiomyocytes provides biochemical and cellular readouts, and in vivo small and large animal models provide information on systemic cardiovascular response. However, there remains a significant gap in these models due to an incomplete recapitulation of adult human cardiovascular physiology, which results in more difficult safety interpretations. Recent efforts in developing in vitro models from engineered heart tissues have demonstrated potential for bridging this gap using human induced pluripotent stem cell-derived cardiomyocytes (hiPSC-CM) in a three-dimensional tissue structure. Here we advance this paradigm by implementing FRESH™ 3D bioprinting to build human cardiac tissues in a medium throughput, well-plate format with controlled tissue architecture, tailored cellular composition, and native-like physiological function, specifically in its adrenergic agonist drug response. To do this, we combined hiPSC-CMs, endothelial cells and fibroblasts in a cellular bioink and FRESH™ 3D bioprinted this mixture in the format of a thin tissue strip stabilized on a tissue fixture. Our results confirmed that FRESH™ 3D bioprinted cardiac tissues could be fabricated directly in a 24-well plate format, were composed of dense and highly aligned hiPSC-CMs at >600 million cells/mL, and within 14 days demonstrated reproducible calcium transients and fast conduction velocity of ∼25 cm/s. Interrogation of these cardiac tissues with the β-adrenergic receptor agonist isoproterenol showed native-like positive chronotropic and inotropic responses, a combination of responses that is not typically observed in 2D monolayer models or standard 3D engineered heart tissue approaches. These results confirm that FRESH™ 3D bioprinted cardiac tissues represents a novel in vitro platform that enables early in vitro pharmacology and toxicology screening.

## INTRODUCTION

Cardiovascular disease is a major burden on the global healthcare system and the leading cause of death worldwide^1^. While there are a range of highly effective cardiovascular drugs for treating high blood pressure, cholesterol, and acute heart failure, there have been very few new drugs approved for treating chronic arrhythmogenic, contractile, or cardiometabolic diseases of the myocardium itself ^2^. Indeed, the number of approved cardiac compounds have stagnated due to high failure rates in clinical trials, leading to the pharmaceutical industry decreasing investment in the development of new cardiac assets, despite the well-recognized healthcare burden^3,4^. These low approval rates and lack of drug pipelines addressing cardiovascular indications can be attributed to the complexity of cardiac physiology and the associated difficulty in appropriately assessing cardiac drug safety and efficacy prior to preclinical and clinical trials^5–7^. Differences between human and animal cardiac physiology limit the quality of insights drawn from preclinical studies, further hampering development efforts^3,8,9^. The FDA Modernization Act 2.0 passed by the United States Congress in 2022^10^ removes the mandate for animal testing to investigate the safety and effectiveness of a drug, thus encouraging drug sponsors to use alternative methods when available. However, for heart disease such in vitro models still need to be improved and validated before they are viable for this application.

The development of human induced pluripotent stem cell derived cardiomyocytes (hiPSC-CMs) has made in vitro culture a promising approach to evaluate drug safety and efficacy, with the potential to supersede certain animal models as the key tool for insight generation in lead optimization prior to human clinical studies^11–17^. Over the past decade two-dimensional (2D) hiPSC-CM monolayer cultures have been adopted with the promise of more physiologically relevant readouts and straightforward integration into conventional well-plate workflows in high-throughput screening^18^. Despite their rapid adoption, growing evidence indicates 2D monolayer hiPSC-CM systems do not fully recapitulate the three-dimensional (3D) structure and function of the adult heart, limiting their value primarily to biochemical and cellular-scale assays^19^. Cardiac spheroids and organoids provide an ability to place hiPSC-CMs in a 3D structure, and this clearly provides some advantages, however the structure and function still recapitulates the embryonic state at best^20^. Engineered heart tissues (EHTs) have been developed to better recreate the 3D structure and function of the heart at the tissue and organ scale, leading to more physiologically data-rich assays^21–30^. Current 3D fabrication approaches for hiPSC-CMs through molding of cell-laden hydrogels, seeding on fiber-based scaffolds and 3D bioprinting have been effective in creating contractile cardiac tissues in a dish. However, there remains a wide range of hiPSC-CM maturation states, tissue structural organization, electrophysiology, contractility, and drug responses.

Here we demonstrate that freeform reversible embedding of suspended hydrogels (FRESH™) 3D bioprinting can be used to fabricate EHTs in a multi-well plate format, taking advantage of the parametric design and robotic control of the additive manufacturing process to control tissue composition, architecture, and reproducibility. FRESH™ 3D bioprinting works by extruding cell-laden bioinks within a gelatin microparticle support bath that provides mechanical support to the bioink while it gels and then can be non-destructively removed by melting at 37°C^31^. We have previously demonstrated that FRESH™ 3D bioprinting can be used to engineer cardiac tissues in a range of complex 3D structures such as ventricle-like constructs and heart tubes with advanced functionality^29,30^. To expand upon this, we created a process that enables EHTs to be FRESH™ 3D printed around custom-designed tissue fixtures within 24-well plates and then performed structural, functional, and pharmacologic assays to demonstrate platform capabilities. Importantly, engineered cardiac tissues have high hiPSC-CM density, uniaxial aligned cells, and exhibit spontaneous contractility. In this report, we establish manufacturing reproducibility, multi-well plate compatibility, and pharmacological chronotropic and inotropic response, which provides a foundation for future applications in toxicology and drug discovery.

## MATERIALS AND METHODS

### Multi-well Plate Design and Fabrication

To create an in vitro array of engineered cardiac tissues, FRESH™ printing was performed within specially designed 24-well plates with integrated tissue fixtures. The tissue fixtures were designed using Solidworks (Dassault Systems) and 3D printed on a Formlabs Form 3B printer (Formlabs) using BioMed Clear (RS-F2-BMCL-01, Formlabs) or BioMed Black resin (RS-F2-BMBL-01, Formlabs). Following printing, the fixtures were cured at 70°C for 60 min, post-processed to remove support material from printing, washed in an isopropanol bath for 5 min, airdried, and then sterilized by autoclaving. The tissue fixtures were then placed within each well of a cell-culture grade 24-well plate (229124, CellTreat).

### FRESH™ Support Bath Preparation

All FRESH™ printing was performed with LifeSupport™, a gelatin microparticle support material prepared according to manufacturer’s directions. Briefly, 1g LifeSupport (FluidForm) was rehydrated in cold (4°C) DMEM/F-12 (11320033, Gibco). Rehydrated LifeSupport was centrifuged, supernatant was removed from the tube, and the compacted LifeSupport was resuspended in DMEM/F-12 containing 1x P/S (P0781, Sigma Aldrich) and 10U Thrombin (91-030, BioPharm Laboratories). Thrombin-supplemented LifeSupport was centrifuged again, and the supernatant was aspirated. A 14G sterile needle was used to transfer the compacted LifeSupport into designated wells of a 24-well plate.

### Cell-laden Bioink Preparation

The cell-laden bioink was prepared as a high-density combination of three cardiac cell types.

Specifically, hiPSC-CMs (iCell Cardiomyocytes^2^ 01434, R1017 Fujifilm Cellular Dynamics), hiPSC-ECs (iCell Endothelial Cells 01434, R1022 Fujifilm Cellular Dynamics), and primary adult ventricular cardiac fibroblasts (C-12375, Promocell). The primary adult ventricular cardiac fibroblasts (CFs) were purchased at low passage, expanded in culture to 70% confluency in fibroblast growth medium (C-23130, Promocell), and cryopreserved for later use. Endothelial cell concentrations were determined based on a recent report^32^. At the time of bioink preparation, all cells were removed from liquid nitrogen storage, thawed, and transferred to media according to manufacturer instructions. Following thaw, the hiPSC-CMs, CFs, and hiPSC-ECs were mixed at the desired cell composition and centrifuged at 200x g for 7 min. The cells were resuspended in 20 mg/mL fibrinogen (9001-32-5, Millipore Sigma) and DMEM/F-12 medium and transferred into a sterile gastight syringe (81230, Hamilton). The gastight syringe was capped, fitted into a custom 3D printed syringe centrifuge adapter, and centrifuged at 200x g for 7 min. The supernatant was aspirated from the syringe and the syringe was fitted with a blunt tip needle, ready for printing. All syringes, needles, adapters, and syringe accessories were sterilized in the autoclave prior to use.

### Design and FRESH™ 3D Bioprinting of Cardiac Tissues

The cardiac tissue was printed within the wells of a 24-well plate with an obround shape suspended between two support posts. The overall dimensions of the cardiac tissue construct as printed were 10 mm length, 2.5 mm width, and 1 mm height, with the tissue strip itself printed with a height of 1 mm and a width of 450 μm. The cardiac tissue was designed in computer-aided design (CAD) software, exported in the STL file format, and then sliced into G-code with a filament diameter of 450 μm and a print speed of 5 mm/s. FRESH™ 3D bioprinting of the cardiac tissue was performed under sterile condition in a biosafety cabinet on a custom-built extrusion 3D bioprinter equipped with a Replistruder 4 syringe pump extruder^33^ For printing, the syringe containing the cell bioink was mounted on the printer and the 24-well plate containing the tissue fixtures and LifeSupport in each well was positioned on the build platform. For each tissue, the needle of the printer was aligned with the print guide on each fixture to ensure accurate placement of the tissue around the fixture posts. After printing, the multi-well plate was transferred to a 37°C incubator for 30 min to melt the LifeSupport and release the cardiac tissues, and then rinsed with DMEM/F12 to remove excess gelatin in the well. The media in each well was then replaced with pen/strep supplemented maintenance medium made from 80% cardiomyocyte medium (M1003, Fujifilm Cellular Dynamics), 20% endothelial cell growth medium (C-22010, Promocell). Cardiac tissues were maintained for up to 4 weeks, with fresh maintenance medium every 2 days.

### Calcium Imaging and Analysis

Calcium imaging was performed to assess the contractility and electrophysiology of the cardiac tissues. For imaging, tissues were incubated at 37°C and 5% O2 for 90 min in Tyrode’s solution containing 5 μM calcium indicator Cal 520 AM (21130, AAT Bioquest) and 0.025% Pluronic F127 (P2443, Sigma Aldrich). Tissues were then transferred into a 50 × 35 mm imaging dish (23000-5035, Kimax) and incubated in Tyrode’s solution for 30 minutes at 37°C for equilibration. Finally, the imaging dish was transferred onto a custom heated stage maintained at 37°C ± 1°C throughout the experiment. High-speed imaging of up to 100 frames per second was performed using a scientific CMOS camera (Prime 95B, Photometrics) mounted on an epifluorescent stereomicroscope (MVX10, Olympus) with a Sola Light Engine and a GFP filter cube. Excitation-contraction decouplers were not used when imaging the entire tissue at 1.25x magnification because there was minimal motion artifact. Custom Python and MATLAB scripts built upon published methods were used to create isochrone maps, calcium traces, quantification of conduction velocity, and calculation of calcium trace metrics^30^. A dynamic tissue mask was generated from each frame and the average pixel intensity of the tissue strip was calculated at each frame and plotted over time. Each data point was then normalized to peak intensity frame at diastole (F/F_0_ -1). In addition, Image J analysis software was used for calcium signal region-of-interest (ROI) analysis based on the criteria that (a) there was minimal motion artifact in that ROI and (b) that the entire ROI contained actively contracting cells.

### Pharmacological Studies

#### 2D Monolayer Studies

The 2D cardiomyocyte monolayer pharmacological studies were conducted using hiPSC-CMs cultured in standard 96-well imaging plates. Briefly, hiPSC-CMs (iCell Cardiomyocytes^2^, Fujifilm Cellular Dynamics) were thawed according to the manufacturer instructions and plated at a density of 30,000 cells/well in 96-well plates (655090, Greiner Bio-One) that had been pre-coated with fibronectin (50 μg/mL) for 3 h at 37°C. CMs were cultured for 48 h in plating media and switched to maintenance media (R1151, Fujifilm Cellular Dynamics). Maintenance media was changed every other day and 24 h prior to the day of the experiment. Cells were assayed 14 days after plating. The hiPSC-CMs were incubated with Codex ACTOne Calcium dye (Codex Biosolutions, Rockville, MD) according to manufacturer instructions for 1 h at 37 °C, 5% CO2. An FDSS/μCell imaging platform heated internally to 37°C (Hamamatsu Ltd., Hamamatsu, Japan) was used as the fluorescence plate reader, with excitation wavelength of 470 nm and the emitted light passed through a 540 nm bandpass filter. A CCD camera simultaneously collected Ca^2+^ transient signals from all 96-wells, at a frame sampling rate of 16 Hz for 1 min. After loading the dye, a baseline recording was taken. Subsequently, isoproterenol was added from compound stocks in DMSO such that the final concentration of DMSO was 0.1%. Final isoproterenol concentrations of 3.7, 11, 33, and 100 nM were applied in triplicate (n=3 wells/concentration) for each batch of iCells tested (total of 3 separate iCell batches). Time-matched vehicle wells (0.1% DMSO) were assayed alongside to normalize for any time- and/or vehicle-dependent effects on the measured parameters of Ca^2+^ transient peak frequency and amplitude. Waveform data from each well were analyzed using the FDSS Wave Analysis software, normalized to baseline of that well, then normalized to time-matched 0.1% DMSO controls, and then averaged. Results are reported as mean (n = 9 wells per concentration).

#### 3D Cardiac Tissue Studies

The pharmacological studies on the FRESH™ printed 3D cardiac tissues were performed after 14 days in culture in Tyrode’s solution using the calcium imaging system already described. Tissues were allowed to equilibrate in the custom heated stage maintained at 37°C ± 1°C and then imaged at baseline and following exposure to isoproterenol (I6504, Sigma Aldrich). For saturation experiments, a single dose of isoproterenol was used. For dose response studies, each dosage of isoproterenol was added sequentially from lowest to highest and each dosage addition was followed by a 10 minute incubation period prior to imaging. Each dosage of isoproterenol was prepared and warmed to 37°C immediately prior to its addition to the tissue to ensure no degradation of drug over time. All drug concentrations were prepared from a 10 mM stock solution of isoproterenol dissolved in DMSO and Tyrode’s solution and were further serially diluted in Tyrode’s solution.

### Immunofluorescence Staining, Imaging, and Analysis

Cardiac tissue strips were fixed and permeabilized overnight at 4°C in 3.7% formaldehyde (252549, Sigma Aldrich) and 0.25% Triton X-100 (T9284, Sigma Aldrich) in 1X phosphate buffered saline (PBS). The tissues were washed and blocked in 4% fetal bovine serum, 0.1% Triton X-100, 50 mM glycine, 1 mM calcium and 1 mM magnesium in 1X PBS overnight at 4°C. The tissues were then washed and stained with 1:100 dilution of mouse anti-sarcomeric α-actinin primary antibody (A7811, Sigma-Aldrich) and 1:100 rabbit anti-CD31 (AB32457, Abcam) for 48 h at 4°C. Following primary antibody staining, tissues were washed and stained for 48 h at 4°C with a 1:100 dilution of goat anti-mouse secondary antibody conjugated to Alexa-Fluor 488 (A28175, Life Technologies), goat anti-rabbit secondary antibody conjugated to Alexa-Fluor 555 (A21428, Life Technologies) at a 1:100 dilution, To-Pro3 at a 1:400 dilution and phalloidin conjugated to Alexa-Fluor 633 (A22284, Life Technologies) at a 1:50 dilution.

Tissues were then washed of excess secondary antibodies. Confocal Z-stacks of tissue strips were acquired using 25x and 63x oil-immersion objective lenses on a Zeiss LSM 980 with Airy Scan. Data was exported to Image J and Imaris 3D segmentation software for processing and visualization. Tissue cell density was calculated in Imaris from the total number of segmented nuclei within the volume of a tissue outlined by actin cytoskeleton staining.

### Statistical Analysis

GraphPad Prism 9 was used for all statistical analysis. A one-tailed, paired T-test was used to evaluate the saturated response of tissue strips to isoproterenol (α = 0.05). The non-linear regression fit function with 4 parameters was used to create dose response curves.

## RESULTS

### Engineering of Cardiac Tissues

FRESH™ 3D bioprinted cardiac tissues were designed and fabricated with several key attributes in mind, specifically, multi-well plate compatibility, biomanufacturing speed and reproducibility, rapid stabilization of tissue function, compatibility with high-quality calcium imaging, and both chronotropic and inotropic response to isoproterenol. Cardiac tissues were FRESH™ printed using a cell-laden bioink prepared with cells directly from cryopreservation, composed of 75% hiPSC-CMs, 10% hiPSC-ECs, and 15% primary adult ventricular human cardiac fibroblasts. This was followed by culture for 14 days to reach a stable state (Figure 1A). The design consisted of a strip of cardiac tissue 450 μm wide and 1 mm high printed in an obround shape around a fixture with two support posts (Figure 1B). Using a cell bioink concentration of 320 million cells/mL, we printed 1.3×10^6^ cells in 4 μL volume of bioink per tissue at the time the cardiac tissue was fabricated. This cell density is approximately an order-of-magnitude greater than that typically used in 3D bioprinting^34,35^. The support fixtures were embedded in LifeSupport within each well and the needle tip of the extruder was aligned to the reference guide for each fixture, which allowed for accurate and precise extrusion and reproducible fabrication of each tissue (Figure 1C). Through the robotic control provided by the 3D bioprinter, manufacturing of a full 24-well plate of cardiac tissues was completed in under 30 minutes (Figure 1D). After printing, the cardiac tissues were cultured for 14 days during which the CFs actively compacted the tissue around the two posts of the support fixture, resulting in dense and highly aligned hiPSC-CMs (Figure 1E). This design also enabled high resolution, high-speed calcium imaging, from which to assess calcium transient and action potential propagation behavior (Figure 1F).

**Figure 1.**
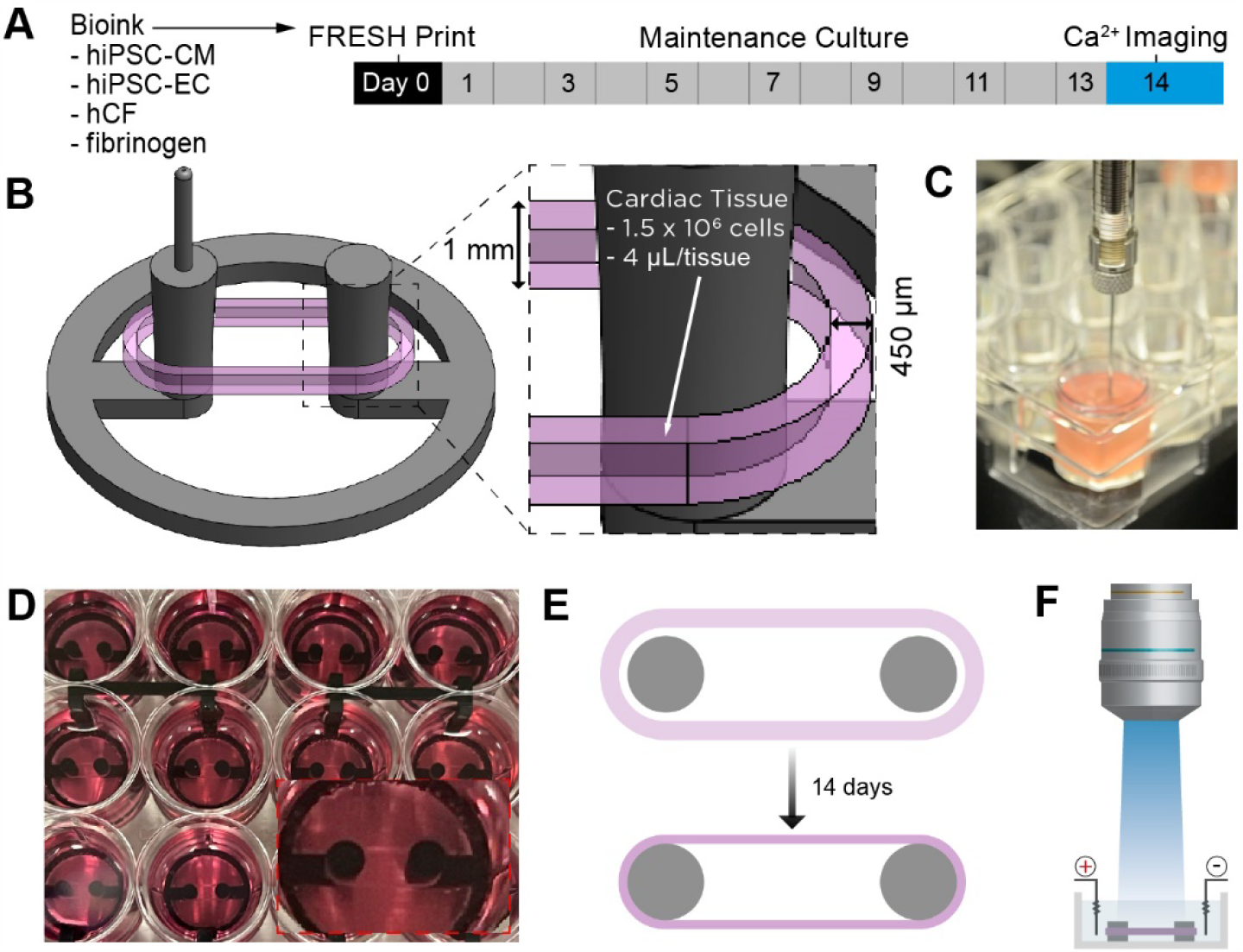
Fabricating FRESH™ printed cardiac tissue strips in a 24-well plate format. (A) Timeline of the fabrication process from FRESH™ printing at day 0 to calcium imaging at day 14. (B) Schematic of the cardiac tissue (pink) designed to be FRESH™ printed around support posts (gray). (C) Image of FRESH™ printing using a cell-laden bioink, printing into a single well. (D) Top-down view of cardiac tissues in each well showing the 24-well plate format, inset shows a cardiac tissue in a single well. (E) Schematic of cardiac tissue compaction around support posts over time in culture. (F) Schematic of the cardiac tissue used for high-resolution, high-speed calcium imaging.

### Structural Characterization of Cardiac Tissues

Following printing, the cardiac tissues underwent a 14-day culture process during which tissues compacted and aligned. Within the first hour post printing, the cardiac tissues began to compact around the pillars of the fixture (Figure 2A). At day 1 post-printing, the cardiac tissue had compacted around the fixturing posts further and rapidly contracting clusters of hiPSC-CMs were dispersed throughout the tissue. By day 7, the cardiac tissues had compacted completely and begun beating synchronously. An additional 7 days of culture, for 14 days total, were required for the cardiac tissue to reach its fully stable shape. At higher magnification the change in cardiac tissue morphology with time could be seen as a compaction and densification of the tissue, reaching a final diameter on the order of ∼150 μm at day 14 (Figure 2B). At day 14 the cardiac tissue also reached a maximum in the calcium transient amplitude, which remained consistent through 28 days in culture (Figure 2C). Whole-mount 3D confocal imaging of the cardiac tissue on day 14 stained for cell nuclei demonstrated the high cell density that was achieved (Figure 2D and Video S1). Cross-sections of the tissue strip in the longitudinal and transverse directions confirmed that cell density was consistent through the entire thickness, and that cell viability was maintained (Figures 2E and 2F). Calculation of nuclei density shows that tissues printed with an initial bioink concentration of ∼320 million cells/mL further compacted to day 14 tissues with cellular density of >600 million cells/cm^3^. This appears to be the highest cell density reported in the literature for an engineered cardiac tissue and is on par with the density reported for neonatal human myocardium^36^. Immunofluorescent staining of sarcomeric α-actinin confirmed that the cardiac tissue contained highly aligned cardiomyocytes in the longitudinal direction of tension from the printing and compaction process (Figure 2G and Figure 2H).

**Figure 2.**
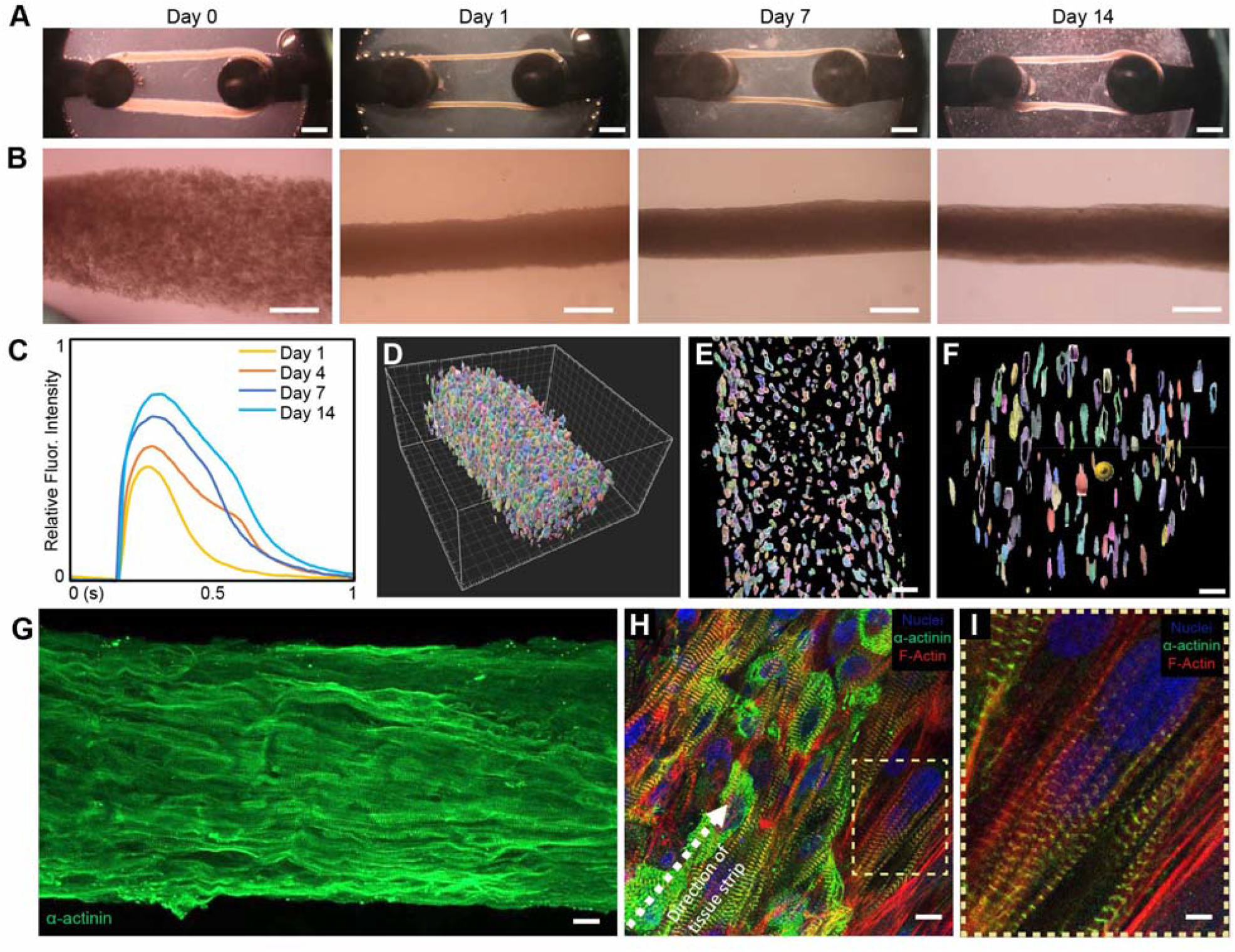
Structural characterization of FRESH™ printed cardiac tissue. (A) Brightfield images of cardiac tissue strips throughout time in culture show compaction and remodeling of tissue strip over time. Scale bars are 1 mm. (B) Higher magnification images of tissues during culture. Scale bars are 100 μm. (C) Calcium transients of cardiac tissues at different time points in culture demonstrating maximum amplitude at 14 days. (D) 3D segmented and color-coded Z-stack of nuclei reveal high cell density throughout the entire tissue, including in (E) longitudinal and (F) transverse cross-sections. Scale bars are 20 μm. (G) Fluorescent image showing uniaxial alignment of sarcomeres with high cell density. Scale bar is 10 μm. (H) High magnification imaging of cardiomyocytes show alignment along length of tissue and (I) well-organized sarcomeric structures. Tissues are stained for nuclei (blue), a-actinin (green), F-actin (red.) Scale bars are 20 μm for (H) and 5 μm for (I).

Further assessment of cardiac subcellular structure deeper within the tissue strip showed well-organized sarcomeres perpendicular to the direction of cellular alignment (Figure 2I).

### Electrophysiology of FRESH™ Printed Cardiac Tissues

The electrophysiology of the FRESH™ printed cardiac tissues was assessed using high-speed and high-resolution calcium imaging. Whole tissue recording of calcium dynamics at day 14 revealed presence of a contractile syncytium with visually observable calcium wave propagation (Figure 3A and Video S2). High magnification imaging of representative regions of interest (ROIs) along the length of tissue displayed comparable calcium trace morphology, indicating the formation of a uniform cardiac tissue substrate with consistent calcium transient properties by day 14 (Figure 3B and Video S3). Conduction velocity in the longitudinal direction of the tissue strips was measured to be 26.9 ±13.5 cm/s (Figure 3C), comparable to the highest values reported for other engineered heart tissues grown under specific electrical stimulation protocols to drive maturation^37^. Consistent with a ventricular phenotype, cardiac tissues developed an automaticity-driven spontaneous beat rate of 45-65 bpm (Figure 3D). When paced under electrical field stimulation, FRESH™ printed cardiac tissues responded as a syncytium and were fully captured at 2 and 3 Hz. Pacing frequencies at or below 1 Hz led to the rise of intermittent contractions between stimulation pulses because of the higher spontaneous beat rate of some tissues.

**Figure 3.**
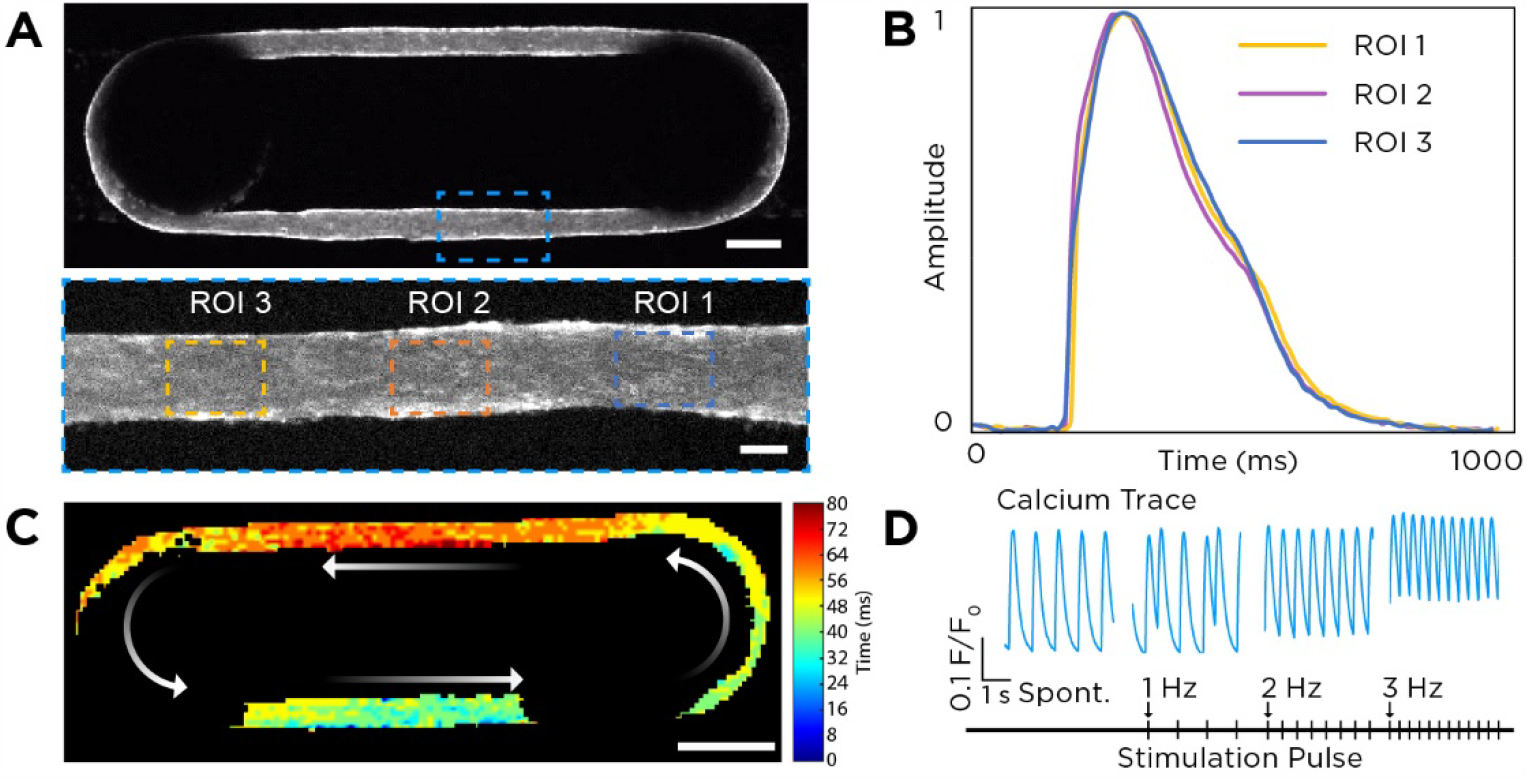
Characterization of FRESH™ printed cardiac tissue calcium handling. (A) Top-down views of a representative cardiac tissue stained with calcium-sensitive dye showing uniform cell distribution. Scale bars are 750 μm (top) and 150 μm (bottom). (B) Plot of calcium transients from the three ROIs in panel (A) showing uniform calcium waveform morphology. (C) Calcium optical mapping of the whole tissue enables observation of the spontaneously propagating action potential and calculation of the conduction velocity. Scale bar is 1 mm. (D) Representative calcium transient for cardiac tissues undergoing spontaneous and electrically stimulated contractions at 1, 2, and 3 Hz.

### FRESH™ Printed Cardiac Tissues Display Positive Chronotropic and Inotropic Response to Isoproterenol

To investigate FRESH™ printed cardiac tissue strips as a platform for pharmacology and toxicology, we assessed response to isoproterenol, a β-adrenergic receptor agonist with well-characterized effects on the adult human heart and in 2D cardiac monolayer cultures. When treated with a saturating level of isoproterenol at 2.5 μM, the FRESH™ printed cardiac tissues responded with ∼40% increase in beat frequency and ∼35% increase in calcium amplitude when compared to untreated baseline (Figure 4A and Video S4). A decrease in calcium cycling time was also observed, marked by both a decrease in calcium transient duration as well as faster relaxation kinetics (Figure 4B). To understand the reproducibility and robustness of this response, we evaluated the isoproterenol-treated calcium dynamics in cardiac tissues FRESH™ printed from an additional commercial hiPSC-CM cell lot (Figure 4C), as well as in cardiac tissues FRESH™ rinted with different compositions of cardiomyocytes, endothelial cells, and cardiac fibroblasts (Figure 4D). Results show that in response to isoproterenol, cardiac tissues fabricated from a different hiPSC-CM line maintained positive chronotropy, and cardiac tissues fabricated with hiPSC-CM and CF but without hiPSC-ECs also maintained a positive chronotropy. Finally, chronotropic dose response to isoproterenol in terms of beat frequency followed the expected curve with the transition from sub-threshold to saturating dose and an EC50 of 1 μM (Figure 4E and Video S5). The FRESH™ printed cardiac tissues also showed a positive relationship between chronotropic and inotropic response as a function isoproterenol dose (Figure 4F). This positive relationship is absent in iCell hiPSC-CMs cultured in 2D monolayers, which show a positive chronotropic response but minimal inotropic response as a function of isoproterenol dose. Thus, the FRESH™ printed cardiac tissues are able to reproduce both the chronotropic and inotropic response of isoproterenol.

**Figure 4.**
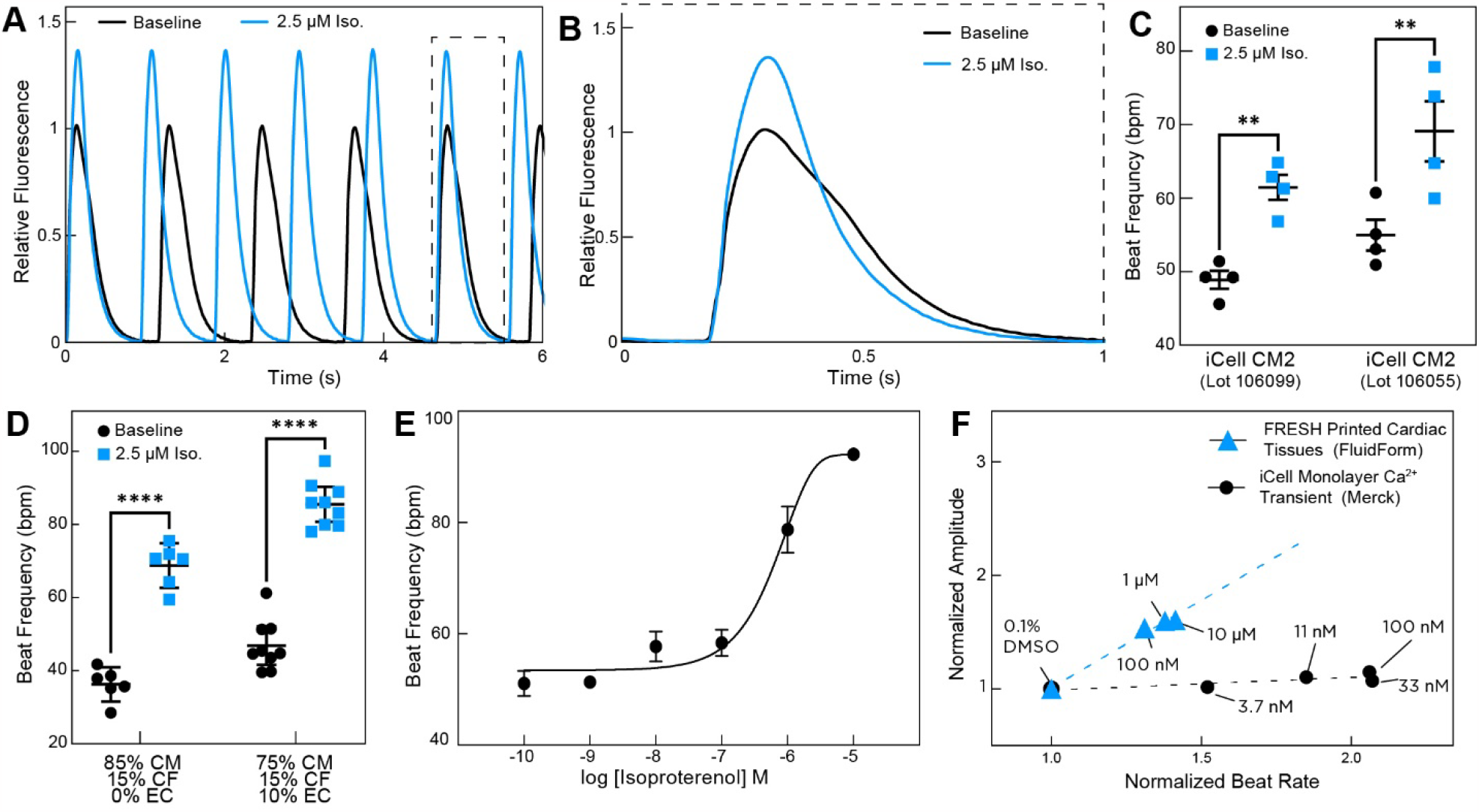
FRESH™ printed cardiac tissues exhibit positive chronotropic and inotropic responses to β-adrenergic receptor agonist isoproterenol. (A) Representative calcium transients from FRESH™ printed cardiac tissue at baseline and exposure to 2.5 μM of isoproterenol. (B) Direct comparison of a single calcium transient for baseline and isoproterenol treated (2.5 μM) shows the characteristic 30-40% increase in peak amplitude. (C) Spontaneous beat frequency in response to 2.5 μM of isoproterenol reveals comparable chronotropic response in cardiac tissues FRESH™ printed with different commercially available hiPSC-CM lines. (D) Spontaneous beat frequency in response to 2.5 μM of isoproterenol reveals comparable chronotropic response in cardiac tissues FRESH™ printed with different cellular compositions, either with two cell types (hiPSC-CMs and CFs), or with three cell types (hiPSC-CMs, hiPSC-ECs, and CFs). (E) Dose response curve of cardiac tissue beat frequency as a function of isoproterenol concentration showing chronotropic response (n = 3). (F) Dose response curves of normalized calcium transient amplitude as a function of normalized beat rate for isoproterenol treated hiPSC-CM 2D monolayers as compared to FRESH™ 3D printed cardiac tissues (n = 4 for FRESH™ printed tissues and n = 9 wells for iCell monolayers).

## DISCUSSION

This study demonstrates well-plate-based manufacturing of cardiac tissues for use in in vitro toxicology and pharmacology. Key to this is the FRESH™ 3D bioprinting process, which enables the use of high-density cell bioinks of >320 million cells/mL that remodel into cardiac tissues with densities of >600 million cells/mL and uniaxially aligned hiPSC-CMs. Except for other work making use of FRESH™ for 3D bioprinting cardiac tissue, we are not aware of any other biofabrication or tissue engineering approaches that can achieve comparable cell densities and uniaxial alignment^30^. We also used commercially available hiPSC-CMs, hiPSC-ECs, and CFs from cryopreservation that were directly combined into the bioink for bioprinting. Not only did this enable us to demonstrate our platform with one of the most widely used hiPSC-CM cell lines in academic and industry research, but it also decoupled the timeline of the cell culture process from the bioprinting process. Overall, this builds off of previous work leveraging FRESH™ bioprinting for the fabrication of functional cardiac tissues^30^ together with other studies working towards utilization of engineered heart tissue for in vitro models in drug discovery^38^. With this platform, we were able to expand upon this previous work by creating a tissue strip designed to be reproducibly manufactured in a multi-well plate format (Figure 1), form cardiac tissue with high density and alignment characteristic of human myocardium (Figure 2), achieve high conduction velocity (Figure 3), and demonstrate positive chronotropic and inotropic response to β adrenoceptor agonist (Figure 4).

The FRESH™ bioprinting technology allowed for the creation of engineered cardiac tissues that overcome several limitations of other cardiac tissue models. Most engineered cardiac tissues are fabricated using cardiomyocytes mixed within a hydrogel matrix at a concentration of 1-30 million cells/mL^11,22,25,39–41^. This relatively low cell concentration is typically driven by the need to have enough hydrogel to achieve adequate mechanical properties when gelled to hold the cardiac tissue together. Here we demonstrated the ability to process and handle a significantly higher cell concentration bioink of (∼320 million cells/mL) through use of FRESH™ printing within a support bath, allowing us to fabricate tissues at cardiomyocyte densities on par with native myocardium^36^. We hypothesize that this native-like cell density in FRESH™ printed cardiac tissues facilitates a high degree of cardiomyocyte alignment and electromechanical coupling, which leads to the observed full function within 14 days of culture and positive chronotropic and inotropic response to isoproterenol (Figure 4E and 4F). This response to β adrenoceptor agonist treatment is typically not observed in other engineered heart tissues or 2D cardiomyocyte monolayers (Figure 4F). Our results demonstrate that FRESH™ 3D bioprinting cardiac tissues can achieve an inotropic response in a hiPSC-CM cell line that fails to do so in 2D culture. Further, FRESH™ printing enabled a design-for-purpose engineering approach, which we leveraged in this study to prioritize design for well-plate compatibility, manufacturing reproducibility, small tissue dimensions, and ease of calcium imaging and analysis. The thinness of the final cardiac tissue strip diameter (∼120 μm) enabled maintenance of tissue viability in a high-density substrate cultured in static conditions. The high aspect ratio of the cardiac tissue strip helped drive cardiomyocyte anisotropy along the longitudinal axis, resulting in the observed conduction velocity of ∼25 cm/s, close to the adult physiologic range of ∼40-60 cm/s^42,43^.

Overall, we were successful in engineering contractile cardiac tissues, however, it is important to note that these tissues do not fully recapitulate adult cardiac function. First, the tissues that we engineered displayed automaticity, a property of fetal cardiac tissues, and common in cardiac tissues engineered from hiPSC-CMs^25,44–48^. Second, our tissues exhibited a flat force-frequency relationship (as quantified by calcium transients), which is different than the positive force-frequency relationship found in adult human myocardium^49,50^. This behavior, together with an absence of post-rest potentiation, suggests immature sarcoplasmic reticulum function compared to the adult heart^51^. Third, while our tissues responded to isoproterenol similar to adult myocardium with positive inotropic and chronotropic response, additional investigation is needed to determine whether this is due to an increase in the I_Ca_ caused by ß-adrenergic receptor activation followed by phosphorylation of the L-type Ca^2+^ channel Ca_V_1.2, or if it is a result of phosphorylation of phospholamban and an increased rate of sarcoplasmic reticulum Ca^2+^ loading^52–54^. Finally, we did not perform ultrastructural analysis to assess whether the hiPSC-CMs contained structural components of mature heart tissue such as the presence of T-tubules. In summary, while we have achieved a number of advances with this platform, there is clearly room for further improvement as the entire field using hiPSC-CMs works towards fully recapitulating the physiology of adult myocardium^16^.

Future work will focus on confirming the capabilities of this platform for in vitro toxicology and pharmacology, both to develop novel disease models for drug discovery^16,17^ as well as to determine the potential to complement or supersede pre-clinical animal testing, when adequately validated. This includes working to improve upon multiple factors that can drive functional maturation, such as the implementation of a dynamic loading regime that can reproduce both preload and afterload. The dynamic engineered heart tissue (dyn-EHT) developed by Bliley et al demonstrates the importance of preload in driving structural and functional maturation of cardiac tissue as compared to most EHT systems that only generate afterload^29^. This dyn-EHT application of preload has also been shown to be critical in driving the emergence of disease phenotypes linked to genetic mutations, such as in arrhythmogenic cardiomyopathy and dilated cardiomyopathy^29,55^. Other factors to implement that can drive cardiac tissue maturation include electrical stimulation during culture^56^, biochemical stimulation such as triiodo-L-thyronine (T3) hormone^57^, and fatty acid metabolism^58–61^. Additional functional readouts are also needed, including contractile force and voltage mapping to quantify the action potential. Cardiac tissues engineered to recapitulate tissue and organ-scale disease states will represent significant next steps to provide clinically translatable data. This includes genetic-based cardiomyopathies (e.g., dilated, hypertrophic, and arrhythmogenic) that can be recreated using patient-derived and/or gene-edited hiPSC-CM lines with monogenic or polygenic mutations^16^. Other forms of heart disease associated with aging also have the potential to be replicated, in the case of FRESH™ 3D bioprinting the reported ability to print collagen^30^ could be used to develop late-stage cardiac fibrosis models through multi-material printing and parametric design based on histology^62^. The tissue design can also be modified to model arrhythmias where irregular electrical signal initiation and propagation can be controlled through multi-material deposition to engineer cardiac substrate heterogeneity in order to potentiate irregularly propagating action potentials.

Taking all of these examples together, it is clear that there are a wide range of approaches to improve engineered cardiac tissue function, and our FRESH™ 3D bioprinted platform has the potential to incorporate many of them.

